# Phosphorylation of pericyte FAK-Y861 affects tumour cell apoptosis and tumour blood vessel regression

**DOI:** 10.1101/2020.06.17.157131

**Authors:** Delphine M. Lees, Louise E. Reynolds, Ana Rita Pedrosa, Marina Roy-Luzarraga, Kairbaan M. Hodivala-Dilke

## Abstract

Focal adhesion kinase (FAK) is a non-receptor tyrosine kinase that is overexpressed in many cancer types and *in vivo* studies have shown that vascular endothelial cell FAK expression and FAK-phosphorylation at tyrosine (Y) 397, and subsequently FAK-Y861, are important in tumour angiogenesis. Pericytes also play a vital role in regulating tumour blood vessel stabilisation, but the involvement of pericyte FAK-Y397 and FAK-Y861 phosphorylation in tumour blood vessels is unknown. Using *PdgfrβCre*+;*FAK*^*WT/WT*^, *PdgfrβCre*+;*FAK*^*Y397F/Y397F*^ and *PdgfrβCre*+;*FAK*^*Y861F/Y861F*^ mice, our data demonstrate that tumour growth, tumour blood vessel density, blood vessel perfusion and pericyte coverage were affected only in late stage tumours in *PdgfrβCre+*;*FAK*^*Y861F/Y861F*^ but not *PdgfrβCre*+;*FAK*^*Y397F/Y397F*^ mice. Further examination indicates a dual role for pericyte FAK-Y861 phosphorylation in the regulation of tumour vessel regression and also in the control of a pericyte derived ‘pericrine’ signals that influence apoptosis in cancer cells. Overall this study identifies the role of pericyte FAK-Y861 in the regulation of tumour vessel regression and tumour growth control and that non-phosphorylatable FAK-Y861F in pericytes reduces tumour growth and blood vessel density.

## Introduction

Angiogenesis is a complex biological process that involves multiple cell types, crosstalk between these cells and responses to different stimuli [1-3]. Interactions between pericytes and endothelial cells play important roles in angiogenesis. Pericyte recruitment to newly forming blood vessels is thought to promote neovessel stabilisation and is an essential step in neovascularisation [4, 5]. Failure to recruit pericytes to blood vessels can affect vascular remodelling, vessel regression and vessel leakage [1, 2, 6]. This has led to pericyte-targeted therapies being developed and a better understanding of the role of pericytes in tumour angiogenesis [7, 8].

It is well documented that focal adhesion kinase (FAK) plays a central role in different aspects of tumour growth and is overexpressed in many types of cancer [9-11]. One of the many roles of endothelial cell FAK, in the promotion of tumour growth, is in the initiation of angiogenesis [12]. FAK regulates growth, survival, migration and invasion through its dual role as a kinase and as a scaffolding protein. FAK-kinase activity results in tyrosine (Y) 397-phosphorylation, which, in turn, allows proteins containing the Src-homology (SH2) domain to bind to FAK, e.g., Src and PI3K. FAK-Src complexing potentiates further FAK phosphorylation at other FAK domains including FAK-Y861 [13, 14]. Most studies have pointed towards essential roles for phosphorylation of FAK-Y397 but much less in known about the requirement of FAK-Y861. We and others have shown previously that endothelial cell (EC) FAK is required for tumour growth since EC FAK loss leads to a reduction in tumour growth, accompanied by a reduction in tumour vascular density [12]. Endothelial specific loss of FAK inhibits brain tumour formation and leads to normalisation of the vasculature [15]. Global constitutive deletion of exon 5, which encodes FAK-Y397, also leads to an embryonic lethal phenotype with vascular permeability defects [16]. More recently, we have shown that non-phosphorylatable mutations of tyrosines 397 and 861 (Y397 and Y861) in endothelial cells have differential effects on tumour angiogenesis [17] and that endothelial cell FAK regulates angiocrine signalling in the control of doxorubicin sensitivity in malignant cells [18]. Whilst FAK has been studied extensively in endothelial cells, the function of FAK in pericytes during tumour growth and angiogenesis has not been explored. Given that pericytes have emerged as an important cell type in the regulation of tumour growth and angiogenesis, we have examined the role FAK in pericytes during pathological angiogenesis. Specifically, we have examined the effect of non-phosphorylatable mutations of tyrosines 397 and 861 (Y397F and Y861F) – FAK residues using pericyte-specific FAK^Y397F/Y397F^ and FAK^Y861F/Y861F^ mice.

In this study, we show that in a subcutaneous Lewis Lung Carcinoma (LLC) tumour model, both angiogenesis and tumour growth are reduced only in mice which have the pericyte-specific FAK-Y861F mutation. This correlates with a significant increase in vessel regression. Furthermore, examination of the secretome and protein expression in FAK-Y861F pericytes highlight an altered ‘pericrine’ signature involving altered cytokines and protein secretion that are involved in cancer cell apoptosis. Thus, pericyte FAK-Y861 plays a role in the control of tumour growth.

## Results

### Generation of pericyte specific FAK mutant mice

We generated a new mouse model that enables us to study the effect of endogenous deletion of Y397 and Y861 of FAK in pericytes, during pathological angiogenesis. In this knock-out/knockin mouse model we initially generated myc-tagged chicken-WT-FAK, or non-phosphorylatable -Y397F or -Y861F mutant FAK constructs (preceded by a STOP sequence flanked by loxP sites) targeted to the Rosa26 (R26) locus. These mice were bred with PC-specific *PdgfrβCre;FAK*^*fl/fl*^ mice to generate mutant FAK-knockin and endogenous FAK-knockout in pericytes under Cre control (**Supplementary Fig. 1a, b**). These mice show no defects in Mendelian ratios, gender distribution or weights and had no obvious adverse phenotype (**Fig. 1a-c**). Pericytes isolated from *PdgfrβCre+;FAK*^*WT/WT*^, *PdgfrβCre+;FAK*^*Y861F/Y861F*^ and *PdgfrβCre+;FAK*^*Y397F/Y397F*^ mice confirmed the presence of the myc-tag indicating chicken FAK knockin, normal levels of total FAK but reduced pY397 in FAK-Y397F pericytes and reduced p-Y861 in FAK-Y861F pericytes. (**Fig. 1 d**).

**Fig 1.**
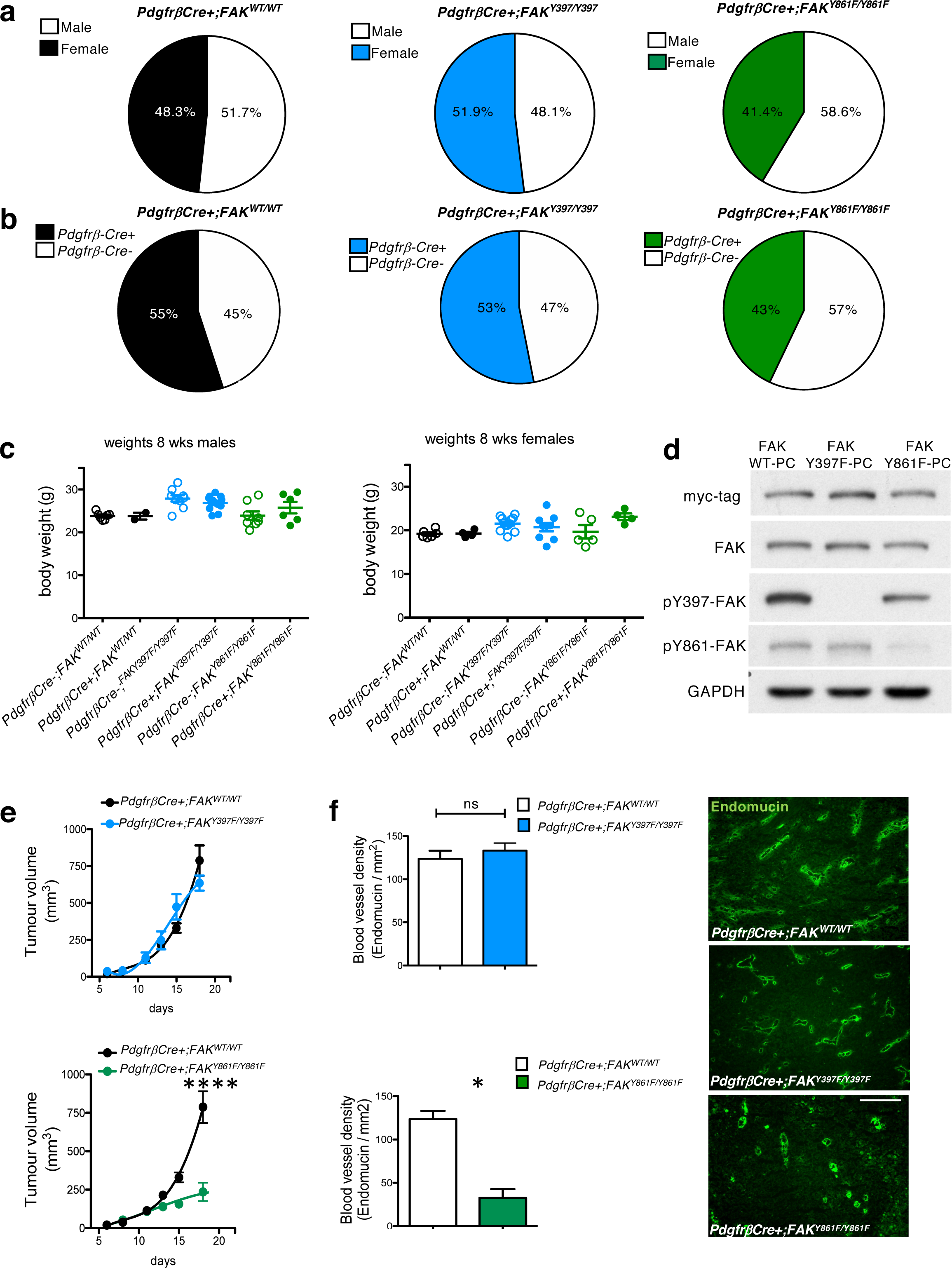
LLC tumour growth and angiogenesis are reduced in PdgfrβCre+;FAK^Y861F/Y861F^ mice. **a** *PdgfrβCre+;FAK*^*WT/WT*^, *PdgfrβCre+;FAK*^*Y397F/Y397F*^ and *PdgfrβCre+;FAK*^*Y861F/Y861F*^ mice were born at normal male;female ratios; **b** Mendelian ratios with **c** similar body weights. Pie chart in **a** represents percentage male:female ratio, in **b** represents % Cre+ and Cre-mice born to each genotype (n=60 mice/genotype). **d** Western blotting of primary brain pericytes isolated from *PdgfrβCre+;FAK*^*WT/WT*^, *PdgfrβCre+;FAK*^*Y397F/Y397F*^ and *PdgfrβCre+;FAK*^*Y861F/Y861F*^ mice confirmed reduced levels of p-Y397 and pY861-FAK in FAKY397F and FAKY861F pericytes respectively; n= 3 cell isolations/genotype. **e** *In vivo* tumour growth was significantly reduced only in *PdgfrβCre+;FAK*^*Y861F/Y861F*^ mice. Graphs represent mean tumour volume ± s.e.m.; n=15 *PdgfrβCre+;FAK*^*WT/WT*^ mice, 14 *PdgfrβCre+;FAK*^*Y397F/Y397F*^ mice and 40 *PdgfrβCre+;FAK*^*Y861F/Y861F*^ mice. ****P<0.0001. Two-sided Mann-Whitney *U* rank sum test. **f** Tumour blood vessel density was significantly reduced only in *PdgfrβCre+;FAK*^*Y861F/Y861F*^ mice. Bar charts represent mean blood vessel density + s.e.m. *P=0.0498; ns, not significant; n= 6 *PdgfrβCre+;FAK*^*WT/WT*^ tumours, 6 *PdgfrβCre+;FAK*^*Y397F/Y397F*^ tumours and 5 *PdgfrβCre+;FAK*^*Y861F/Y861F*^ tumours. Two-sided Student’s *t*-test. Representative endomucin stained LLC tumour sections are shown for each genotype. Scale bar, 50 μm.

### Tumour growth and blood vessel density are reduced in *PdgfrβCre+;FAK*^*Y861F/Y861F*^ but not *PdgfrβCre+;FAK*^*Y397F/Y397F*^ mice

To examine the effects of pericyte FAK-Y397F and Y861F mutations on tumour growth and angiogenesis, *PdgfrβCre+;FAK*^*WT/WT*^ control mice *PdgfrβCre+;FAK*^*Y861F/Y861F*^ and *PdgfrβCre+;FAK*^*Y397F/Y397F*^ mice were injected subcutaneously with Lewis Lung Carcinoma cells (LLC). *In vivo* tumour growth was reduced in *PdgfrβCre+;FAK*^*Y861F/Y861F*^, but not *PdgfrβCre+;FAK*^*Y397F/Y397F*^ mice. Furthermore, the *PdgfrβCre+;FAK*^*Y861F/Y861F*^ but not *PdgfrβCre+;FAK*^*Y397F/Y397F*^ mice had significantly reduced blood vessel density (as determined by the number of endomucin-positive vessels per mm^2^ of age-matched, size-matched tumours) compared with *PdgfrβCre+;FAK*^*WT/WT*^ control mice (**Fig. 1e, f**). In addition to the above, blood vessel perfusion (determined by the percentage of endomucin positive blood vessels that were also positive for PE-PECAM antibody after ante-mortem perfusion) and pericyte coverage (determined by the percentage of vessels with NG2-postive mural cell association) were both reduced in *PdgfrβCre+;FAK*^*Y861F/Y861F*^ but not *PdgfrβCre+;FAK*^*Y397F/Y397F*^ mice (**Fig. 2 a, b**). Loss of blood vessels can result from partial or persistent regression of vascular endothelial cells. Blood vessel regression is associated with endothelial cell loss and pericyte dropout leaving behind empty collagen IV basement membrane sleeves [19, 20]. Thus, vessel regression can be determined by the presence of endothelial cell-negative, collagen IV basement membrane blood vessel sleeves. Collagen IV / endomucin double-immunostaining of day 14-21 tumours (when the rate of tumour growth in *PdgfrβCre+;FAK*^*Y861F/Y861F*^ mice slows down) from *PdgfrβCre+;FAK*^*WT/WT*^, *PdgfrβCre+;FAK*^*Y397F/Y397F*^ and *PdgfrβCre+;FAK*^*Y861F/Y861F*^ mice showed a significant increase in endomucin-negative / collagen IV-positive blood vessel sleeves in tumours from *PdgfrβCre+;FAK*^*Y861F/Y861F*^ compared with tumours from *PdgfrβCre+;FAK*^*WT/WT*^, *PdgfrβCre+;FAK*^*Y397F/Y397F*^ mice (**Fig. 2c)**. These results correlate with the reduction in tumour growth and suggest that tumour blood vessel regression is central to the tumour phenotype in *PdgfrβCre+;FAK*^*Y861F/Y861F*^ mice.

**Fig 2.**
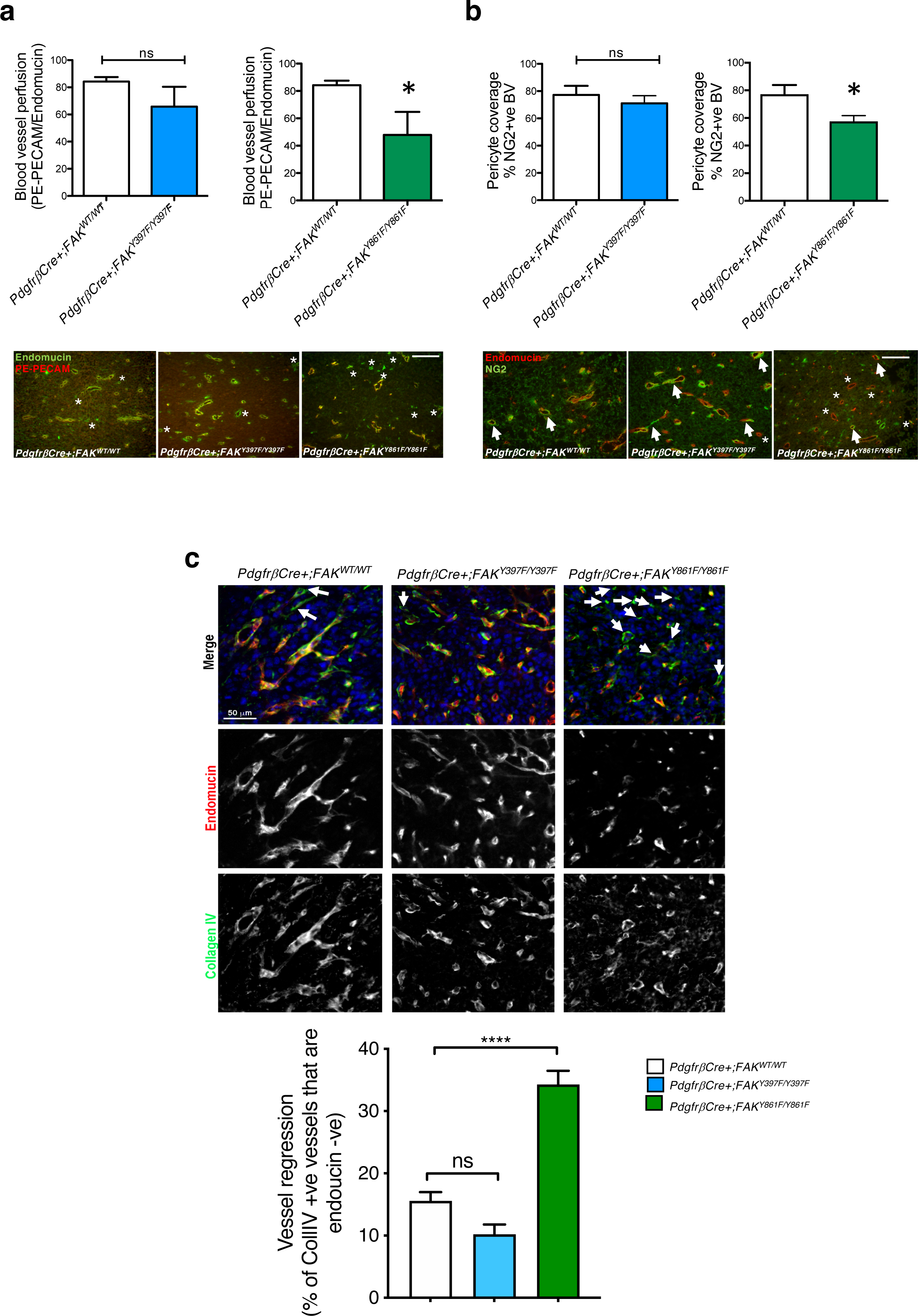
Blood vessel perfusion and pericyte coverage are reduced in PdgfrβCre+;FAK^Y861F/Y861F^ mice. **a** PE-PECAM perfused LLC tumours grown in *PdgfrβCre+;FAK*^*WT/WT*^, *PdgfrβCre+;FAK*^*Y397F/Y397F*^ and *PdgfrβCre+;FAK*^*Y861F/Y861F*^ mice showed a significant reduction in functional tumour blood vessels in *PdgfrβCre+;FAK*^*Y861F/Y861F*^ mice. Bar charts represent mean blood vessel perfusion + s.e.m. *P=0.032; ns, not significant; n= 6 *PdgfrβCre+;FAK*^*WT/WT*^ tumours, 6 *PdgfrβCre+;FAK*^*Y397F/Y397F*^ tumours and 5 *PdgfrβCre+;FAK*^*Y861F/Y861F*^ tumours. Two-sided Student’s *t*-test. Representative endomucin stained and PE-PECAM perfused LLC tumour sections are shown for each genotype. *asterisks*, non-perfused blood vessels. **b** Pericyte coverage of tumour blood vessels was also reduced in these mice. Bar charts represent mean pericyte coverage + s.e.m. *P=0.03, ns, not significant; n= 6 *PdgfrβCre+;FAK*^*WT/WT*^ tumours, 6 *PdgfrβCre+;FAK*^*Y397F/Y397F*^ tumours and 5 *PdgfrβCre+;FAK*^*Y861F/Y861F*^ tumours. Two-sided Student’s *t*-test. Representative double stained endomucin and NG2 LLC tumour sections are shown for each genotype. *Arrows*, NG2+ endomucin+ blood vessels; *asterisks*, NG2-endomucin+ blood vessels. Scale bar in **a** and **b**, 50 μm. **c** Day 14-21 tumours from *PdgfrβCre+;FAK*^*WT/WT*^, *PdgfrβCre+;FAK*^*Y397F/Y397F*^ and *PdgfrβCre+;FAK*^*Y861F/Y861F*^ mice were immunostained with Collagen IV and endomucin to identify empty basement membrane sheaths. Bar chart shows vessel regression (% of Coll IV+ vessels that are endomucin -ve) + s.e.m.; ****P< 0.0001; ns, not significant; n=6 *PdgfrβCre+;FAK*^*WT/WT*^ mice, 8 *PdgfrβCre+;FAK*^*Y397F/Y397F*^ mice and 6 *PdgfrβCre+;FAK*^*Y861F/Y861F*^ mice. Two-way ANOVA. Representative images show Collagen IV and endomucin stained blood vessels from tumours from all genotypes. *arrows*, Collagen IV+ endomucin –ve blood vessels. Scale bar, 50 μm.

### Increased tumour necrosis in early stage tumours of *PdgfrβCre+;FAK*^*Y861F/Y861F*^ mice

Examination of early stage tumours was undertaken to determine at what stage, during growth of the tumour, angiogenesis was being affected. In early stage tumours (day 12 post tumour cell inoculation) LLC tumour size, blood vessel density and blood vessel perfusion were similar between *PdgfrβCre+;FAK*^*WT/WT*^ and *PdgfrβCre+;FAK*^*Y861F/Y861F*^ mice (**Fig. 3a-c**), suggesting that loss of blood vessels occur at a later stage of tumour growth. Indeed to examine whether FAK-Y861F PCs could directly affect the initial stages of microvessel sprouting, aortic rings from *PdgfrβCre+;FAK*^*Y861F/Y861F*^ and *PdgfrβCre+;FAK*^*WT/WT*^ micewere embedded in collagen and stimulated, or not, with VEGF (30ng/ml). VEGF treatment significantly increased angiogenic sprouting in both genotypes to the same extent suggesting that mutation of FAK-Y861F in pericytes is not sufficient to directly affect endothelial cell sprouting in a tumour free environment (**Supplementary Fig. 2**).

**Fig 3.**
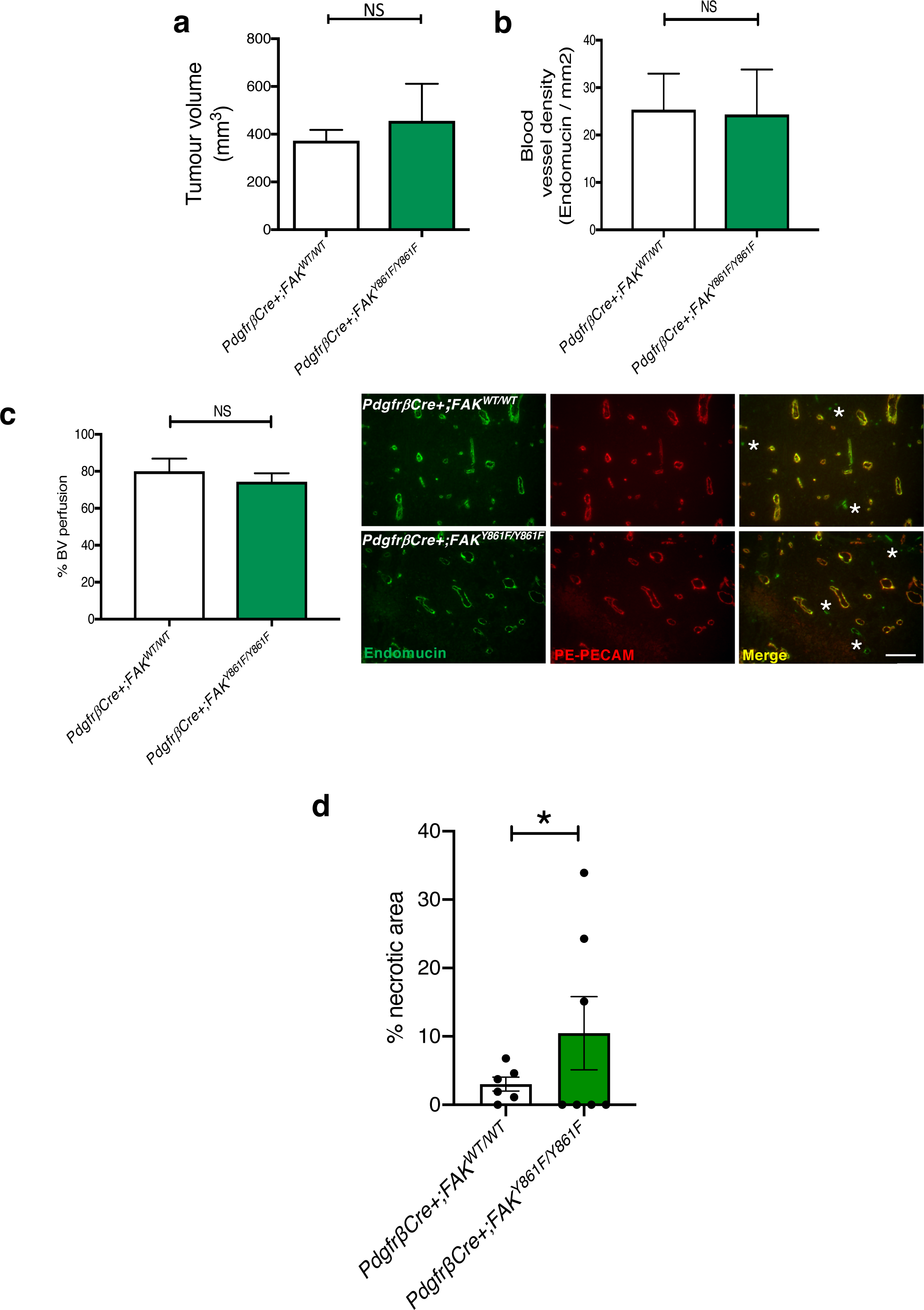
Early stage tumour growth is similar between PdgfrβCre+;FAK^WT/WT^ and PdgfrβCre+;FAK^Y861F/Y861F^ mice Early (day 12): **a** Tumour size, **b** Blood vessel density, and **c** Blood vessel perfusion were similar between *PdgfrβCre+;FAK*^*WT/WT*^ and *PdgfrβCre+;FAK*^*Y861F/Y861F*^ mice. Bar charts represent mean + s.e.m.; ns, not significant. Student’s *t*-test. For **a**, n= 8 *PdgfrβCre+;FAK*^*WT/WT*^ mice, *5 PdgfrβCre+;FAK*^*Y861F/Y861F*^ mice. For **b** and **c** n= 8 *PdgfrβCre+;FAK*^*WT/WT*^ tumours, 5 *PdgfrβCre+;FAK*^*Y861F/Y861F*^ tumours. Representative images showing blood vessel perfusion in LLC tumours from *PdgfrβCre+;FAK*^*WT/WT*^ and *PdgfrβCre+;FAK*^*Y861F/Y861F*^ mice. Scale bar, 50 μm. *Asterisk*s, non-perfused blood vessels. **d** Tumour necrosis was increased in *PdgfrβCre+;FAK* ^*Y861F/Y861F*^. Bar chart shows % necrotic area in tumours from *PdgfrβCre+;FAK*^*WT/WT*^ and *PdgfrβCre+;FAK* ^*Y861F/Y861F*^. *P<0.05 Wilcoxon test; n=6 tumours/genotype.

Examination of early stage tumours, using H&E stained sections, revealed that tumours from *PdgfrβCre+;FAK*^*Y861F/Y861F*^ mice had a significant increase in tumour necrosis compared with *PdgfrβCre+;FAK*^*WT/WT*^ control mice (**Fig. 3d**). These results suggest that tumour necrosis precedes the onset of vessel regression and the reduced late stage tumour blood vessel density and tumour growth in *PdgfrβCre+;FAK*^*Y861F/Y861F*^ mice.

### FAK-Y861F pericyte secretome enhances LLC tumour cell apoptosis

Paracrine or juxtacrine signals from endothelial cells, angiocrine signals, have been implicated in the control of tumour growth where endothelial cell derived signals directly affect tumour cells by altering cytokine profiles [18, 21, 22]. Much less is known about the possible role of pericytes in this type of signalling, that we coin ‘pericrine’ signalling. Thus, we turned our attention to the possible effects of FAK-Y861F pericytes on tumour cell apoptosis. Using R&D protein profiler cytokine arrays, lysates from FAK-Y861F pericytes showed a decrease in levels of thrombospondin, MCP-1, proliferin, TIMP-1 and sICAM/CD54 together with an increase in IGFBP-2, endostatin TNF-alpha SDF1 and ADAMTS-1 (**Fig. 4a**). Given that *in vitro* MCP-1 has been shown to promote mural cell recruitment [23] and *in vivo* pharmacological inhibition of MCP-1 reduces tumour growth and macrophage recruitment resulting in increased tumour necrosis [24] whilst loss of MCP-1 delays mammary tumourigenesis [25], we examined the effect of treating LLC with MCP-1 after exposure to Y861F conditioned medium (CM) and asked if MCP-1 could be involved in controlling tumour cell apoptosis. CM from FAK-WT and FAK-Y861F pericytes were incubated with cultured LLC cells and apoptosis quantified by TUNEL staining. Y861F CM caused a significant increase in LLC apoptosis compared with LLCs incubated with WT CM. Indeed the pro-apoptotic phenotype of Y861F CM was rescued by the addition of exogenous recombinant MCP-1 to the pericyte CM (**Fig. 4b**), suggesting that the reduction of MCP-1 in Y861F pericytes is at least partially responsible for the pro-apoptotic phenotype in tumours grown in *PdgfrβCre+;FAK*^*Y861F/Y861F*^ (Y861F) mice.

**Fig 4.**
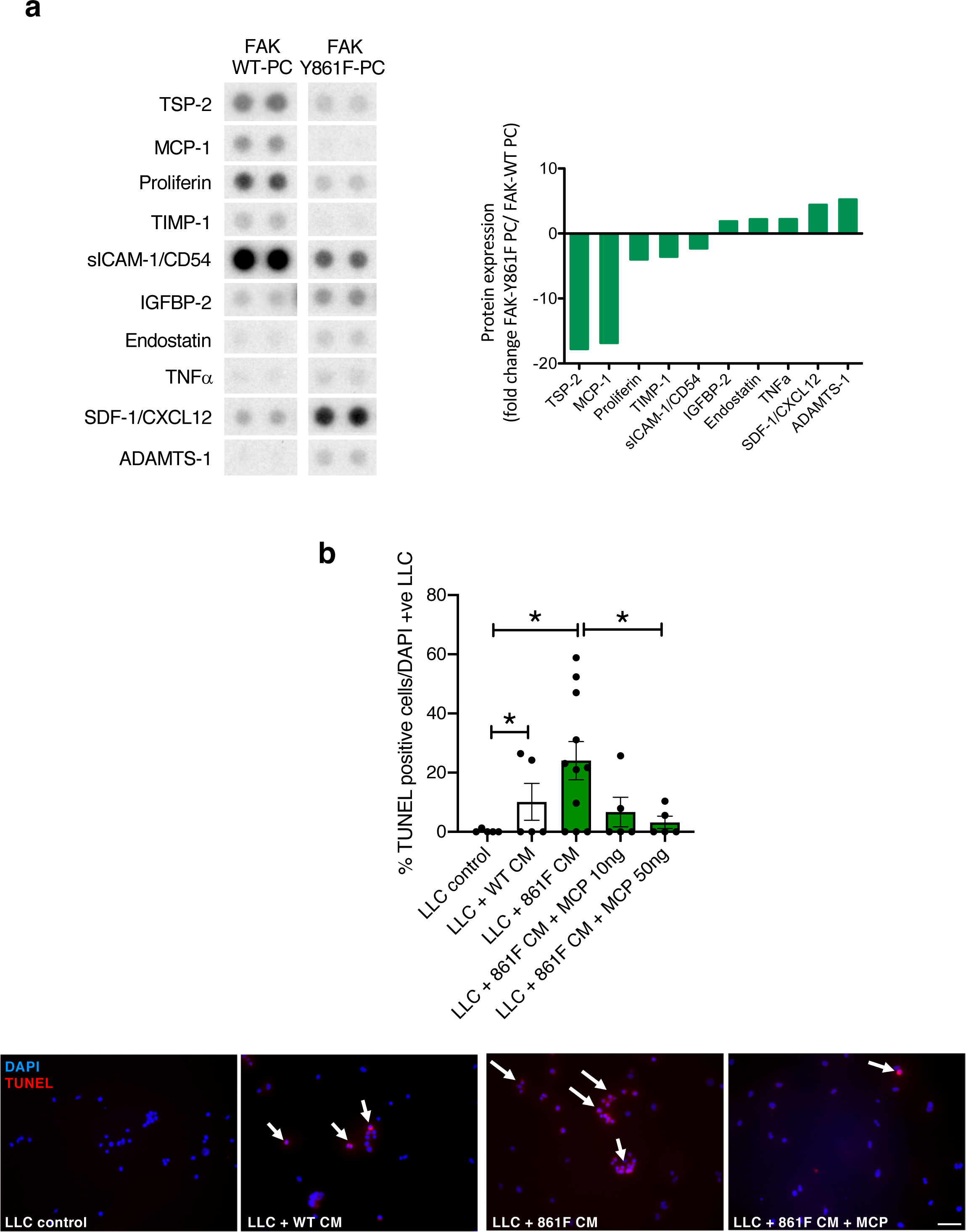
FAK-Y861F pericytes induce apoptosis in LLC tumour cells. **a** R7D proteome cytokine array using lysates from FAK WT and FAK-Y861F pericytes. Representative dots of differentially expressed proteins are given. Bar chart shows mean fold change in protein expression. N= 2 dots from 1 experiment. **b** Lewis lung carcinoma (LLC) cells incubated with conditioned medium (CM) from pericytes plus or minus recombinant MCP-1. Treatment with FAK-Y861F pericyte CM increased LLC apoptosis, compared with CM from FAK-WT pericytes. This effect was rescued upon treatment with MCP-1 (10 and 50 ng/ml). Bar chart represents % TUNEL positive cells + s.e.m. Representative images show effect of CM and MCP-1 on LLC cells. *Arrows*, TUNEL+ cells. *P<0.05. One-way ANOVA. N= 3 technical repeats. Scale bar, 500 μm.

These results imply a direct regulation of tumour cells through “pericrine” signalling - a direct effect of PCs on tumour cells which has an effect on vascular disruption and ultimately tumour growth.

## Discussion

Our data demonstrate that FAK-Y861F pericytes decrease tumour burden in part by directly affecting tumour cell apoptosis, and call for further consideration of the role of tumour pericytes in the direct control of tumour growth, in addition to their effects on vessel stabilisation.

FAK is upregulated in many cancer types and is currently being targeted as a potential anti-cancer agent because of its essential roles in tumour growth and angiogenesis [12, 26-28]. Our previous work identified opposing roles for FAK-Y397 and FAK-Y861 phosphorylation in tumour endothelial cells where endothelial cell FAK-Y397 but not FAK-Y861 reduced tumour growth and angiogenesis by altering signaling pathways downstream of FAK [17, 29]. Pericytes are thought to be essential in stabilising the tumour vasculature and thus attention has been given to the potential of targeting pericytes to induce vascular disruption in cancer control [30]. However, recent conflicting studies have emerged challenging this idea [31] and suggest possible additional roles for pericytes in regulating tumour growth. In contrast to the effect of FAK-Y397F mutation in endothelial cells [17], our mouse models show that phosphorylation of FAK-Y397 in pericytes is not a prerequisite for the control of LLC tumour growth or angiogenesis. The reasons for this apparent discrepancy likely reflect a different requirement for FAK-Y397-phosphorylation in these two cells types in tumour blood vessels. FAK-Y861F mutation in pericytes is associated with reduced tumour growth and angiogenesis that is preceded by an increased tendency for tumour cell necrosis and subsequent blood vessel regression. Since tumour necrosis has been demonstrated to have vascular disrupting effects [32, 33] it is tempting to speculate that the early stage enhanced tumour necrosis may initiate vascular disruption. Additionally, given that the Y861F mutation reduces pericyte association with tumour blood vessels, it is likely that vessel stability and maturation is compromised and this reduced pericyte coverage is also likely to be responsible for the decrease in the numbers of functional blood vessels and reduced tumour growth.

Studies have shown that vascular endothelial cells can control tumour growth via angiocrine signals, including cytokines and chemokines [18, 34]. In our study, the protein signature of FAK-Y861F pericytes is a combination of reduced levels of TSP-2, MCP-1, proliferin, TIMP-1 and sICAM/CD54 together with an increase in IGFBP-2, endostatin, TNF-alpha, SDF1 and ADAMTS-1. This molecular signature associates with pro-apoptotic effects on tumour cells but apparently has little effect on VEGF-stimulated angiogenic sprouting in aortic ring assays in a tumour-free context. The lack of an effect on VEGF-induced aortic ring sprouting suggests that the effect of FAK-Y861F pericyte enhancement in tumour cell apoptosis may be a contributing factor to subsequent vessel regression in a tumour environment. The anti-angiogenic effect of TSP-2 is blocked by VEGF thus providing a possible explanation of why VEGF-stimulated angiogenesis is not affected aortic ring assays from *PdgfrβCre+;FAK*^*Y861F/Y861F*^ mice where the decrease in TSP-2 might otherwise have increased angiogenesis [35]. The pro-tumoural effects of MCP-1, proliferin, TIMP-1 and soluble ICAM-1 [36-39] whilst the anti-cancer effects of IGFBP-2, endostatin and ADAMTS-1 [40-42] have been published and suggest that this secretome signature may well provide a combination of effects on cancer growth. On the contrary, SDF1 has cancer promoting effects [43] but the fold increase in *PdgfrβCre+;FAK*^*Y861F/Y861F*^ cells is much smaller than the downregulation of TIMP-1 and MCP-1, suggesting that the cumulative effect of the secretome is towards an anti-cancer effect. Indeed MCP-1 addition to conditioned medium from FAK-Y861F pericytes was able to rescue the pro-apoptotic effect of this cocktail on tumour cells. The molecular mechanism by which FAK-Y861 affects this pericrine signature is the subject of the future study. Our data support the notion that pericytes are not simply vessel supporting cells, but that via FAK-Y861, can regulate tumour growth via pericyte derived signals directly affecting tumour cell apoptosis.

## Methods

### Mice

To examine the effect of pericyte-specific FAK mutations *in vivo*, we developed *PdgfrβCre;FAK*^*W/WT*^ mice (FAK WT), *PdgfrβCre;FAK*^*Y397F/Y397F*^ (Y397F) mice and *PdgfrβCre;FAK*^*Y861F/Y861F*^ (Y861F) mice [44]. All experiments were approved by United Kingdom Home Office Regulations. For animals bred in-house, health screens (quarterly) were conducted in accordance with FELASA guidelines for health monitoring of rodent colonies, to confirm their free status of known pathogens in accordance with FELASA screens. No clinical signs were detected. Animals were housed in groups of 4-6 mice per individually ventilated cage in a 12 h light dark cycle (06:30-18:30 light; 18:30-06:30 dark), with controlled room temperature (21 ± 1 °C) and relative humidity (40-60 %). The cages contained 1-1.5 cm layer of animal bedding, and with environmental enrichment including cardboard Box-tunnel and crinkled paper nesting material. Animals had access to food and water ad libitum.

### Tumour growth

Male and female mice aged 12-16 weeks were injected subcutaneously with 0.5×10^6^ Lewis lung carcinoma cells (LLC, ATCC) into the flanks. Tumour dimensions were measured over time and tumour growth was determined using the formula: length x width^2^ x 0.52. When tumours reached the maximum legal size allowed, mice were killed, tumour sizes measured and tumour samples were either snap-frozen, fixed in 4 % paraformaldehyde (PFA) or fixed in 4 % PFA/sucrose for histological analysis. For early tumour growth studies, tumours were harvested at day 12 post tumour cell injection.

### Blood vessel density

Five μm frozen tumour sections were air-dried for 10 min, washed once in PBS, fixed in acetone for 10 min at - 20°C, washed in PBS three times and then blocked with 5 % normal goat serum for 30 min at room temperature. After blocking, sections were incubated with primary antibodies overnight at 4°C. Primary antibodies used were directed against endomucin (clone V.7C7; Santa Cruz; sc-65495, 1:100). Sections were then washed with PBS and incubated with Alexa-Fluor^®^-conjugated secondary antibody (1:100, Invitrogen) for 45 minutes at room temperature before mounting the slides with Prolong^®^ Gold anti-fade reagent (Invitrogen, P36934). Tumour blood vessels were counted across entire midline sections, and the numbers were expressed as vessels/mm^2^. For examination of blood vessel regression, tumours were fixed with 4 % PFA/sucrose then frozen in OCT.

### Blood vessel perfusion

For analysis of the % of functional tumour vessels, 100 μl PE-CD31 antibody (clone 390, neat; Biolegend, 102407) was injected via the tail vein 10 min prior to culling mice. Tumours were dissected immediately, snap frozen and sectioned. Frozen sections were then immunostained for endomucin, as described above. To calculate the % number of functional vessels, the number of PE-CD31-positive blood vessels was divided by the total number of endomucin-positive blood vessels.

### Pericyte coverage

Frozen tumour sections were double immunostained as described in “Blood vessel density” using the pericyte-specific antibody NG2 (AB5320; Millipore, 1:100) and endomucin. The percentage of endomucin-positive vessels with NG2-positive cells associated was calculated.

### Blood vessel regression/Collagen IV staining

Frozen tumour sections were air dried for 30min at room temperature, permeabilised for 3 min with PBS + 0.5 % TritonX-10 then blocked with 5% BSA/PBS for 1 hr at room temperature. Sections were then incubated overnight at 4°C with Collagen IV (Abcam; 1:100 dilution, ab6586) and endomucin (clone V.7C7, Santa Cruz; 1:100) antibodies. The following day, sections were washed three times with PBS and incubated for 1 hr at room temperature with AlexaFluor conjugated secondary antibodies (1:100). Finally, sections were washed twice in PBS and once in distilled water then mounted (Prolong Gold with DAPI) with a glass slide and images were acquired using a confocal spinning disk microscope and sCMOS confocal camera (Nikon). Image analysis was performed using ImageJ software by making maximum intensity projections of z stacks and the numbers of vessels counted manually.

### Tumour necrosis

Early stage LLC tumours were fixed in 4% formaldehyde and bisected. Tumour sections were H&E stained, scanned using a Panoramic scanner and the area of necrosis (as identified by acellular regions of tumour tissue) quantified with ImageJ software.

### MCP-1 treatment of LLC with pericyte conditioned medium

LLC were plated on coverslips at a density 5 x10^4^ cells in DMEM + 10% FCS. At the same time WT and 861F pericytes were grown to 50-60% confluency and conditioned medium (CM) removed. CM was centrifuged to remove cell debris and added to LLCs after removal of DMEM and two PBS washes. Recombinant mouse MCP-1 (Biotechne, MAB479) was added to the CM at either 10 or 50 ng/ml. Cells were incubated with CM for 24 or 48h after which cells were stained to detect DNA fragmentation in apoptosis using the BrdU-Red DNA (TUNEL) kit (Abcam, ab66110) following the manufacturer’s instructions. Within 3 hours of staining the cells were analysed for BrdU using a Zeiss microscope and Axiovision software. The percentage of TUNEL positive cells was calculated by counting the total number of cell nuclei and the number of nuclei that were TUNEL positive.

### Primary cell cultures

Primary mouse brain pericytes were isolated from the mice, cultured and characterised as previously described [45, 46]. Briefly, brains were removed from mice, minced, digested for 1 hour in 0.1 % collagenase, centrifuged in the presence of 22 % BSA, and cultured in endothelial cell growth media (pMLEC) with the medium changed every 3 days. On reaching confluency, cultures were harvested with trypsin and passaged. During the first two passages, pericyte cultures were grown in pMLEC, and on the third passage they were grown in pericyte medium (PM; ScienCell Research Laboratories) containing 2 % FBS and antibiotics. Tissue culture plates for all experiments were coated with a mixture of collagen (30 μg/ml), gelatin (0.1 %) and fibronectin (10 μg/ml).

### Aortic ring assay

Aortic rings were isolated from all mouse genotypes as previously described [47].

### Angiogenesis and cytokine arrays

Pericyte angiogenesis and cytokine profiles were compared using the angiogenesis array (ARY015, R&D Biosystems) and cytokine array (ARY006, R&D Biosystems). Briefly, cell lysates were prepared as follows: sample buffer was added to the cell culture, the cells were scraped and transferred into a 1.5 ml Eppendorf tube. After sonication, samples were adjusted to the array conditions and mixed with a Detection Antibody Cocktail as indicated by the manufacturer’s instructions. Lysates were incubated overnight at 4°C on dot-blot membranes. Membranes were washed, incubated with streptavidin-HRP for 30 min at RT, washed again and ECL was applied to the membrane to reveal the dots. Quantification of dot intensity was performed using ImageJ software.

### Western blot analysis

Primary brain pericytes were grown to 70-80 % confluency then lysed in RIPA buffer. 15-30 μg protein was run on 8 % polyacrylamide gels then transferred to nitrocellulose membranes. Membranes were probed with primary antibody overnight at 4°C. Myc-tag (Cell Signaling, clone 9B11, 2276, 1:1000), total FAK (Cell Signaling, 3258, 1:1000), phospho-397 FAK (Invitrogen, 44-624G, 1:1000), phospho-861 FAK (Invitrogen, 44-626G, 1:1000), PDGFRβ (Cell Signaling, clone 28E10, 3169, 1:1000), endomucin (V7.C7, Santa Cruz, 1:1000). The anti-HSC70 (Santa Cruz, clone B-6, sc-7298) or GAPDH (Millipore, MAB374) antibody, for loading controls, were used at 1:5000 dilution. Densitometric readings of band intensities were obtained using the ImageJ™ software.

### Statistical analysis

Statistical significance was calculated using Prism 8 software and *P*<0.05 was considered statistically significant, unless otherwise indicated. For tumour growth statistics, non-parametric two-sided Mann-Whitney *U* rank sum test was performed to compare tumour volumes each day. One-way ANOVA was performed for the TUNEL and aortic ring assay, two-way ANOVA for the blood vessel regression study. Wilcoxon test was performed for tumour necrosis.

## Acknowledgments

We would like to thank Julie Holdsworth and Bruce Williams from Barts Cancer Institute (QMUL, UK) for their expert technical assistance and Professor Ralf Adams (Max Planck Institute for Molecular Biomedicine, Münster, Germany) for the kind gift of the *PdgfrβCre* mice.

## Declarations

### Ethics approval

All procedures were approved by our local animal ethics committee, Queen Mary University of London, and were executed in accordance with United Kingdom Home Office regulations.

### Consent to participate

Not applicable

### Consent for publication

### Availability of data and material

The data that support the findings of this study are available from the corresponding author upon request.

### Code availability

Not applicable

### Authors’ Contributions

DL designed and executed the experiments. LER wrote the paper, executed the experiments, analysed data and gave scientific advice. RP performed collagen IV staining and analysis. MR-L analysed tumour necrosis slides. KMH-D devised and supervised the project and wrote the paper.

## SUPPLEMENTARY INFORMATION

**Supplementary Fig 1 *Generation of PdgfrβCre+;FAK***^***WT/WT***^, ***PdgfrβCre+;FAK***^***Y397F/Y397F***^ ***and PdgfrβCre+;FAK***^***Y861F/Y861F***^ ***mice.* a** Schematic representation of WT FAK, the non-phosphorylatable tyrosine 397 (Y397F) and tyrosine 861 (Y861F) mutations. **b** Generation of *PdgfrβCre+;FAK*^*WT/WT*^, *PdgfrβCre+;FAK*^*Y397F/Y397F*^ and *PdgfrβCre+;FAK*^*Y861F/Y861F*^ mice.

**Supplementary Fig 2 *VEGF-stimulated microvessel sprouting using PdgfrβCre+;FAK***^***WT/WT***^ ***and PdgfrβCre+;FAK***^***Y861F/Y861F***^ ***aortic rings.*** Graph shows mean number of sprouts/ring ± s.e.m. Each data point is the number of sprouts for an individual ring. N=24 *PdgfrβCre+;FAK*^*WT/WT*^ aortic rings/treatment; n=21 *PdgfrβCre+;FAK*^*Y861F/Y861F*^ aortic rings/treatment. ****P<0.0001, One-way ANOVA; ns, not significant.

