## Supplemental File for "Phosphorylation of pericyte FAK-Y861 affects tumour cell apoptosis and tumour blood vessel regression"

**SUPPLEMENTARY INFORMATION**

**Supplementary Fig 1** ***Generation of PdgfrβCre+;FAK$^{WT/WT}$, PdgfrβCre+;FAK$^{Y397F/Y397F}$ and PdgfrβCre+;FAK$^{Y861F/Y861F}$ mice.*** **a** Schematic representation of WT FAK, the non-phosphorylatable tyrosine 397 (Y397F) and tyrosine 861 (Y861F) mutations. **b** Generation of *PdgfrβCre+;FAK$^{WT/WT}$*, *PdgfrβCre+;FAK$^{Y397F/Y397F}$* and *PdgfrβCre+;FAK$^{Y861F/Y861F}$* mice.

**Supplementary Fig 2** ***VEGF-stimulated microvessel sprouting using PdgfrβCre+;FAK$^{WT/WT}$ and PdgfrβCre+;FAK$^{Y861F/Y861F}$ aortic rings.*** Graph shows mean number of sprouts/ring ± s.e.m. Each data point is the number of sprouts for an individual ring. N=24 *PdgfrβCre+;FAK$^{WT/WT}$* aortic rings/treatment; n=21 *PdgfrβCre+;FAK$^{Y861F/Y861F}$* aortic rings/treatment. ****P<0.0001, One-way ANOVA; ns, not significant.

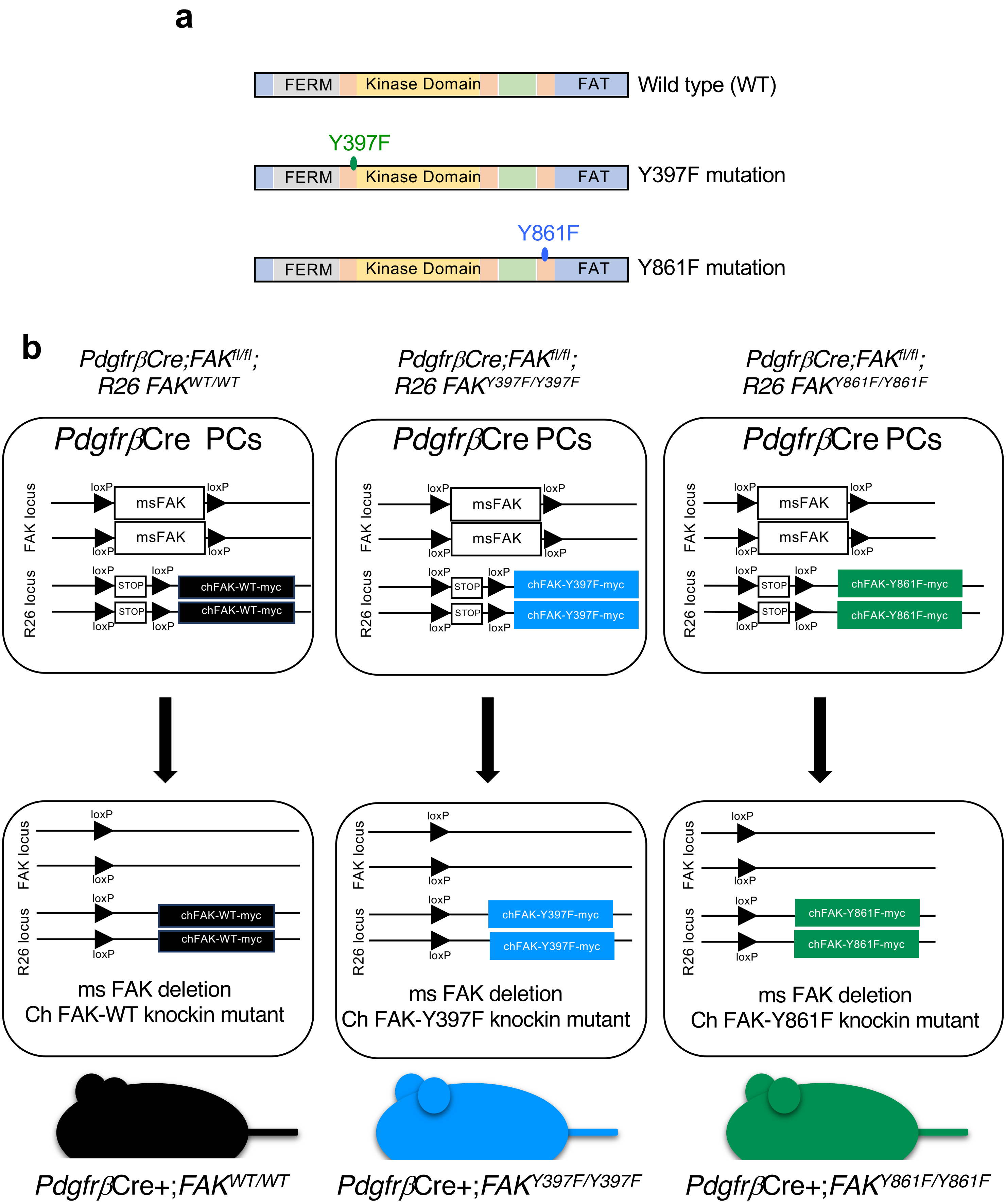
a
FERM
Kinase Domain
FAT
Wild type (WT)
Y397F
Y397F mutation
Y861F
Y861F mutation
b
PdgfrβCre;FAKfl/fl; R26 FAKWT/WT
PdgfrβCre;FAKfl/fl; R26 FAKY397F/Y397F
PdgfrβCre;FAKfl/fl; R26 FAKY861F/Y861F
PdgfrβCre PCs
FAK locus
R26 locus
loxP
msFAK
STOP
chFAK-WT-myc
chFAK-Y397F-myc
chFAK-Y861F-myc
ms FAK deletion
Ch FAK-WT knockin mutant
Ch FAK-Y397F knockin mutant
Ch FAK-Y861F knockin mutant
PdgfrβCre+;FAKWT/WT
PdgfrβCre+;FAKY397F/Y397F
PdgfrβCre+;FAKY861F/Y861F

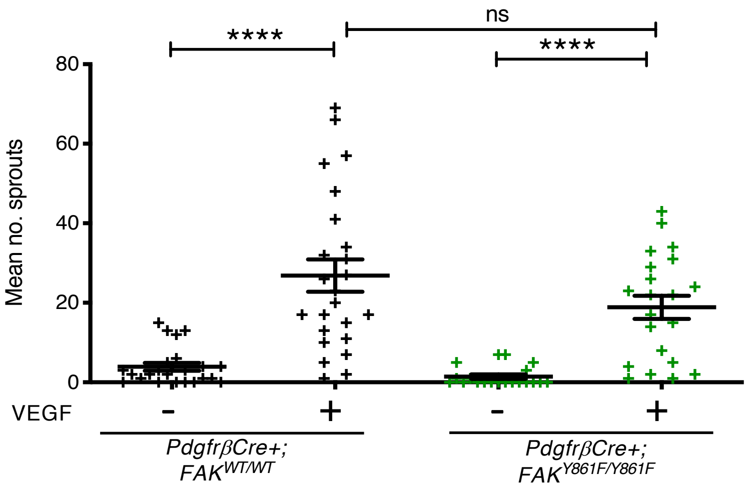

ns
****
****
80
60
40
20
0
Mean no. sprouts
VEGF
-
+
-
+
PdgfrβCre+;
FAK WT/WT
PdgfrβCre+;
FAK Y861F/Y861F
